# Morphofunctional changes at the active zone during synaptic vesicle exocytosis

**DOI:** 10.1101/2022.03.07.483217

**Authors:** Julika Radecke, Raphaela Seeger, Anna Kádková, Ulrike Laugks, Amin Khosrozadeh, Kenneth N. Goldie, Vladan Lučić, Jakob B. Sørensen, Benoît Zuber

## Abstract

The fusion of synaptic vesicles (SVs) with the plasma membrane (PM) proceeds through intermediate steps that remain poorly resolved. Additionally, the effect of persistent high or low exocytosis activity on intermediate steps remains unknown. Through time-resolved cryo-electron tomography, we ordered events into a sequence. Following stimulation, proximal tethered SVs rapidly form additional tethers with the PM. Simultaneously, fusion initiation occurs by membrane curvature (‘buckling’) of the SV and PM. It is followed by the formation of a fusion pore, and the collapse of SV membrane. At this time, membrane-proximal, but not membrane-distal, vesicles lose their interconnections, allowing them to move towards the PM. Two mutations of SNAP-25 that arrests or disinhibit spontaneous release, respectively, both caused a loss of interconnectors, while the disinhibiting mutant also caused a loss of membrane proximal multiple-tethered SVs. Overall, tether formation and connector dissolution is triggered by stimulation and respond to the spontaneous fusion rate. These morphological observations likely correspond to the transition of SVs from one functional pool to another.

## Introduction

For normal brain function such as movement coordination or memory formation, communication between neurons is essential. In the central nervous system, neurons communicate through the release of neurotransmitters at synapses. This process relies on synaptic vesicle (SV) exocytosis, i.e. the fusion of neurotransmitter-filled SVs with the plasma membrane (PM). SV exocytosis involves a sequence of steps [1,2]. The vesicle is docked to the active zone (AZ) PM and the exocytosis machinery goes through a maturation process, termed priming, after which the SV is ready to fuse. These SVs form the readily releasable pool (RRP). Finally, a calcium influx triggers fusion of the SV with the PM. Docked SVs are defined as those in very close proximity or direct contact with the PM as observed by electron microscopy (EM), whereas priming refers to SV ability to undergo exocytosis immediately upon stimulation. Whether every docked SV is also primed has been debated [1,3,4]. A high-pressure freezing/freeze-substitution EM study of synapses has indicated that vesicles which are in direct contact with the PM, i.e. docked, are also primed and belong to the RRP and that this situation occurs downstream of vesicle tethering [4]. From a molecular perspective, priming involves several proteins, including the SNARE complex (SNAP-25, syntaxin-1, and synaptobrevin-2), Munc13, Munc18, synaptotagmin-1, and complexin [2,5]. All three SNAREs form a highly stable tight four-helix bundle, known as trans-SNARE complex. The surfaces of the SV and the PM, are both negatively charged and therefore tend to repulse each other. The formation of the trans-SNARE complex counteracts this repulsion and brings the SV and the PM in high proximity [6]. Evidence has suggested that the SNARE complex is only partially zipped in primed vesicles [7]. Furthermore, various studies have suggested that the formation of at least three SNARE complexes provides the necessary energy for a vesicle to become fusion-competent [8,9,10]. Yet in the absence of cytoplasmic Ca^2+^, minimal spontaneous exocytosis takes place. When the presynaptic terminal gets depolarized by an action potential, Ca^2+^ flows in the cytoplasm and binds to synaptotagmin-1, which is localized at the SV surface. Upon Ca^2+^ binding, synaptotagmin-1 was proposed to insert between the head groups of the PM anionic phospholipids and trigger membrane curvature and destabilization, leading first to hemifusion and subsequently to fusion [11]. Interestingly, much of the trans-SNARE bundle surface is negatively charged. This contributes to the electrostatic barrier that minimizes spontaneous fusion. Synaptotagmin-1 can then act as an electrostatic switch that triggers exocytosis. Introducing negatively charged side chains by site-directed mutagenesis reduces the rate of spontaneous and evoked exocytosis, whereas introducing more positive side chains enhances the rate of spontaneous exocytosis and depletes the RRP. [12]

Cryo-electron tomography (cryo-ET), which preserves samples to atomic resolution, revealed that under resting conditions, no SV is in direct contact with the PM and the majority of AZ-proximal SVs are connected to the PM by a variable number of short tethers [13,14]. The observed gap between the SV and the PM is consistent with the model of an electrostatic barrier formed by the negative charges of the SV, the PM, and the trans-SNARE bundle [12]. In synaptosomes treated with hypertonic-sucrose solution, which depletes the RRP, the majority of tethered vesicles had only 1 or 2 tethers [13,15,16]. This observation suggested that the RRP consists of SV, which are linked to the PM by 3 or more tethers. The RRP, as identified by morphological criteria, only represents a minority of AZ-proximal vesicles. This is in agreement with previous reports. In one of them, the term pre-primed pool was used for the few vesicles (~1 vesicle at hippocampal synapses) that are rapidly released and another publication showed that the RRP is made up of only 10-20% of SVs located on the AZ (equal to ~1 vesicle on hippocampal synapses) [17,18]. The ensemble of proximal vesicles that are not in the RRP have been termed non-RRP and presumably belong to the recycling pool that releases more slowly [13,19]. Farther away from the AZ, partially intermixed with the recycling pool, is the reserve pool containing vesicles that only release upon high frequency stimulation. Vesicles in the reserve pool are tightly clustered and well inter-connected by structures that were termed connectors [13,19]. It should be noted, that the molecular nature of connectors is not known and is possibly heterogenous. Synapsin has been proposed as a molecular constituent. However, since the deletion of all forms of synapsin does not lead to the complete absence of connectors, it is clear that not all connectors contain synapsin [20,21]. The second row of SVs near the active zone (45-75 nm from AZ), immediately after the proximal vesicles (<45nm from AZ), is called the intermediate region. Resting state intermediate SVs are less densely packed and also less connected than proximal SVs [14]. This suggests that, after exocytosis of RRP SVs, intermediate SVs could be rapidly recruited in the RRP by diffusion [22]. Synaptic activity enhances the mobility of a fraction of SVs, whereas it induces synapsin dissociation from SVs in a synapsin phosphorylation-dependent manner [23,24]. The same mobility enhancement can be achieved through inhibition of synapsin dephosphorylation, which leads to synapsin dissociation from SVs, or by knocking out all three synapsin forms [25,26,27]. Interestingly, ribbon synapses do not express synapsin and show higher SV mobility than conventional synapses [28]. It is therefore conceivable that inter-SV connectors restrain SV diffusion and that synaptic activity influences the level of inter-SV connectivity and thereby their mobility.

To investigate this hypothesis and to better understand the impact of depolarization and synaptic activity on SV tethering, we designed two sets of cryo-ET experiments. On the one hand, we compared the morphology of wild-type rat synaptosomes in resting state and a few milliseconds after depolarization. On the other hand, to study the consequences of increased or decreased spontaneous synaptic activity, we imaged synapses in mouse neuronal culture expressing either wild-type SNAP-25, a more positively charged SNAP-25 mutant (4K mutant), or a more negatively charged mutant of SNAP-25 (4E mutant) [12] The more positively charged SNAP-25 mutant, which is constitutively active, showed no triple-tethered SV [12]. This confirmed the morphological definition of the RRP. Our experiments revealed an immediate formation of additional tethers between proximal RRP vesicles and the PM after depolarization. Shortly after exocytosis the level of inter-SV connectivity was decreased among SVs situated in a 25 to 75-nm distance range from the AZ PM. Altogether, our results indicate a regulation through connectors of SV mobility and their recruitment at the AZ PM.

## Results

To analyze the morphological changes occurring in the presynapse shortly after stimulation, we pursued a time-resolved cryo-electron tomography approach similar to the one introduced by Heuser, Reese, and colleagues [29]. Whereas Heuser et al. used an electrical stimulus, we chose to trigger exocytosis by spraying a depolarizing solution containing 52-mM KCl onto the specimen a few milliseconds before freezing for two reasons. Firstly, cryo-ET requires samples to be plunge-frozen directly on an EM grid, which is not compatible with electrical stimulation. Secondly, this method allows to catch synapses at delays between stimulation and freezing lower than a millisecond, as explained below. The delays achieved here are shorter than those attained by electrical stimulation given the uncertainty on the exact time of freezing. The solution was sprayed with an atomizer and droplets hit the EM grid a few milliseconds before freezing. The spray-mixing plunge freezing setup was custom built based on a system introduced by Berriman and Unwin [30]. The spray droplet size was optimized by cutting a 1-ml pipet tip to a diameter matching an EM grid (3 mm) and fixed to the atomizer glass outlet to disperse the spray (Figure 1A1). Furthermore, to achieve the shortest possible delay between spraying and freezing, the nozzle was set 1-2 mm above the liquid ethane container. This generated many small spray droplets spread throughout the grid (Figure 1A2-A4, Supplementary Figure S1). Even if sprayed droplets were well distributed throughout the grid, not all synaptosomes were in contact with exocytosis-triggering KCl solution. Synaptosomes located on the landing spot of a droplet were stimulated instantly and therefore were frozen for a delay equal to the time between the grid crossing the spray and hitting the cryogen (typically set at 7 or 35 ms; Supplementary Figure S1B). However, synaptosomes situated at a distance from a landing spot were only stimulated when the KCl concentration rose due to diffusion reached a threshold triggering voltage-gated calcium chanel opening (Supplementary Figure S1C and D). Through this process, we were able to trap stimulated synapses at the very earliest stages of exocytosis. Given the very low throughput of cryo-electron tomography, we followed a correlative light and electron microscopy approach. By cryo-fluorescence microscopy, we identified areas where synaptosomes fluorescently labeled by calcein blue and spray droplets labeled by fluoresceine were colocalized. Additionally, phase contrast imaging enabled quality control of the frozen EM grid with respect to ice contamination and ice cracks, as shown previously [31]. 9 control and 9 stimulated synaptosome tomograms were analyzed (Supplementary Figure S2A and B, Supplementary Movie S1, Supplementary Table S1). We restricted our analysis to synaptosomes that possessed a smooth PM, free of signs of rupturing and that had a mitochondrion, as we considered these factors essential for synaptosome function.

In addition, we manipulated the electrostatic state of the SNARE complex through mutated SNAP-25 protein introduced using lentiviral vectors into primary SNAP-25 knockout neurons grown on EM grids [12] (Figure 1B1-B4). The “4E” mutation contains four glutamic acid substitutions, whereas the “4K” contains four lysine substitutions; all mutations are placed in the second SNARE-motif of SNAP-25 and were shown to decrease and increase the rate of spontaneous miniature release, respectively [12]. Optimization of Primary neurons culturing conditions allowed us to establish a protocol, which provides functional synapses thin enough for direct imaging by cryo-ET. Astrocytes were added to 12 well plates and were grown for 2 days. After 2 days, the medium was exchanged to a medium that favors neuronal growth and impedes astrocyte growth. At the same time a droplet of the neuronal suspension was added onto a flame sterilized EM grid and incubated for 30 min at 37 °C, hereafter the grids were placed into the 12 well plates containing the astrocytes. Neurons were grown for 10-14 days until plunge freezing and were then analyzed at a Titan Krios by cryo-ET (Supplementary Figure S2C and D, Supplementary Figure S3, Supplementary Movie S2, Supplementary table S2). Thereby, we could image chronically overactive or inactive synapses and relate presynaptic architectural modifications to different functional states.

**Figure 1:**
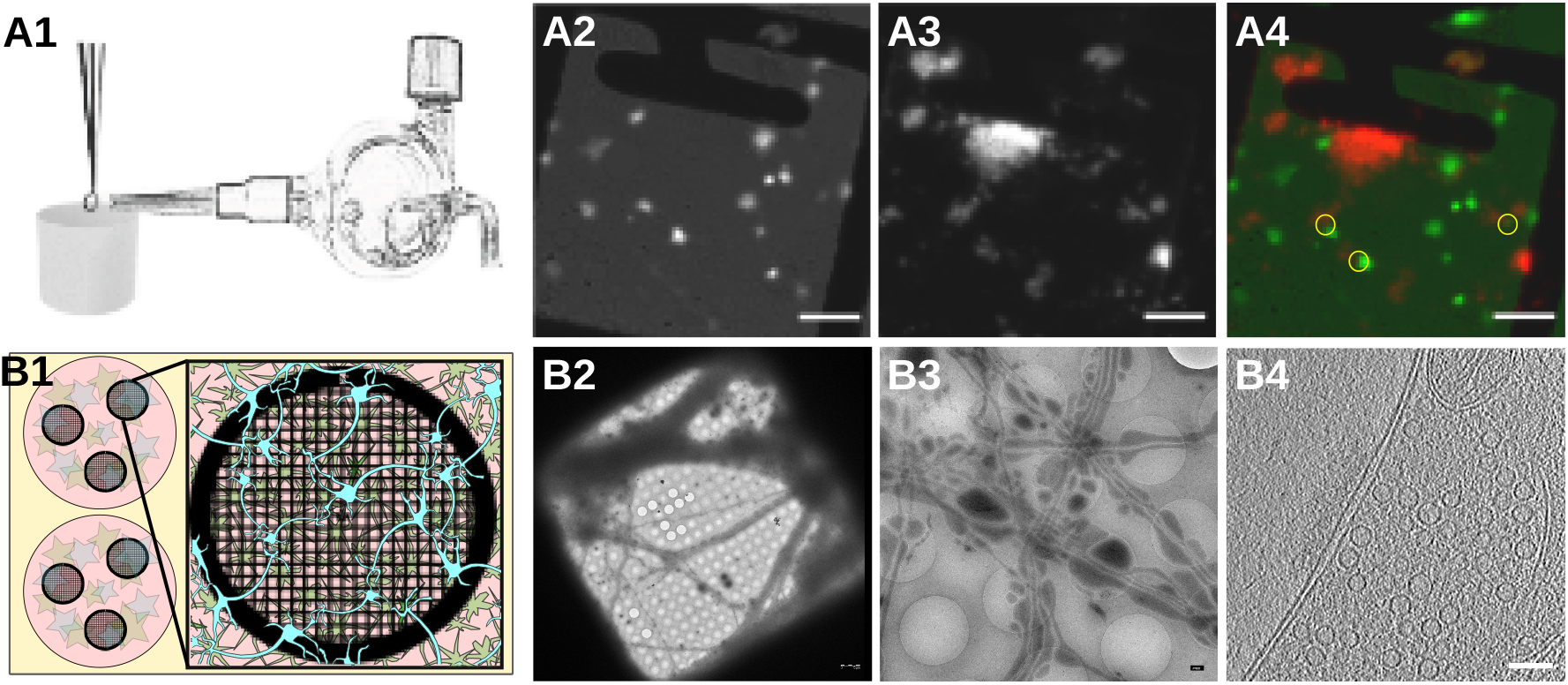
Experimental models. A1) Glass atomizer used to disperse depolarizing solution on the EM grid milliseconds before the grid is vitrified. A2) Spray droplets imaged with the GFP filter set. A3) Synaptosomes imaged with the DAPI filter set. A4) Overlay of spray droplets (green) and synaptosomes (red). Scale bars, 20 μm. B1) Schematic drawing of a 6-well petridish depicting astrocytes (pink) growing at the bottom of the petridish below EM grids (black round grid overlaying the astrocytes) with neurons (blue) growing on top of the grids. B2) Gridsquare overview with neurons growing over it; scale bar = 5 μm. B3) Medium magnification overview of neurons growing over R2/1 holes; scale bar = 500 nm. B4) One slice of a tomogram depicting the synapse and its respective postsynapse; scale bar = 100 nm.

### Increased membrane curvature at the onset of exocytosis

We analyzed the morphology of SVs fusing with the AZ PM. As explained above, synaptosomes of a single grid have not all been stimulated for the same duration (Supplementary Figure S1)· The time interval between triggering exocytosis and freezing ranged between 0 ms and the interval between spray droplets hitting the grid and freezing, which was comprised between 7 and 35 ms depending on the experiments (see [30]). This offered the unique possibility to observe SV exocytosis events immediately after their initiation, and even before membranes have started to mix.

Synaptosomes from both control and sprayed grids were thoroughly analyzed for signs of exocytosis, which consisted of morphological changes of the AZ PM and the tethered SV occurring upon stimulation, which are described hereafter. These signs were only detected in synaptosomes from sprayed grids. We analysed non-sprayed tomograms not only acquired specifically for this study but also from past studies and from hundreds of SVs at the active zone we found no sign of exocytosis. Thereafter the snapshots of exocytosis are presented in the most parsimonious chronological order. Upon stimulation, both the vesicle membrane and the PM were slightly bent towards each other (Figure 2B1-B3; orange arrows). These structures, which have previously been reported in liposomes but not in synapses, have been referred to as membrane curvature events [11]. Control synaptosomes (i.e. not sprayed) on the other hand, had a straight PM, and no SV membrane was buckled (Figure 2A). Following membrane bending, we observed contacts between vesicles and the PM bilayer where both membranes lose their clear contours (Figure 2C1 & C2; pink arrows). This was followed by further transitioning states prior to and during pore opening (Figure 2D-F; blue arrows). In the next observed fusion state, the vesicle was wide open (Figure 2G), followed by almost completely collapsed vesicles where only a small bump on the PM remained visible (Figure 2H). These structures were not observed in any of the non-sprayed control datasets.

**Figure 2:**
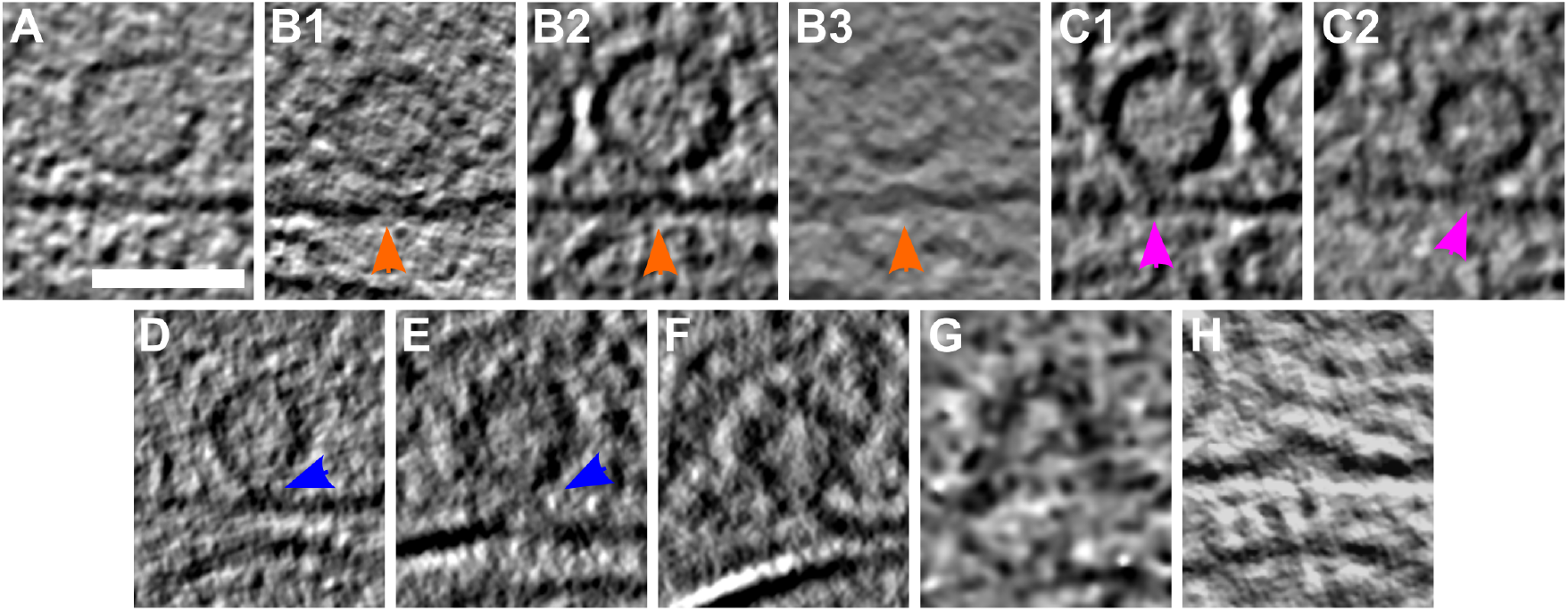
SV exocytosis morphology. Tomographic slice of non-stimulated (A) and stimulated rat synaptosomes (B-H). A) Image of a 2.2-nm thick tomographic slice showing a non-stimulated with SVs at the AZ and a straight PM. B1) Membrane curvature event, 2.2-nm thick tomographic slice. B2) Membrane curvature event, 6.5-nm thick tomographic slice. B3) Membrane curvature event, 2.24 nm thick tomographic slice. Orange arrows showing membrane curvature event. C1,C2) Lipid perturbations of PM and SV, 22-nm thick tomographic slices. The space between SV and PM is denser than in the non-stimulated synaptosomes (see pink arrow). D-F) Vesicles with a pore opening that might be on the way to full collapse fusion, 33-nm thick tomographic slice thickness: 22 nm (D), 30.8 (E), 33 nm (F). G) Wide pore opening, most likely on the way to full collapse fusion, 2.2-nm tomographic slice. H) Remaining bump at the end of full collapse fusion, 11-nm thick tomographic slice. Scale bar, 50 nm.

Stimulated synaptosome datasets were divided into early and late fusion stages, respectively, based on the morphology of SV and AZ PM. Synapses showing membrane bending and direct lipid contact between SV and PM without an open pore were classified as early fusion (Figure 2B-E). Those with an open pore or a remaining small bump of a fully collapsed vesicle were classified as late fusion (Figure 2F-H).

### Synaptic vesicle distribution is impacted by synaptic activity

Non-sprayed rat synaptosomes as well as WT-SNAP-25 mouse cultured neuron synapses showed typical SV distribution, as observed in previous cryo-ET studies (Figure 3)[13]. Vesicle occupancy in WT-SNAP-25 synapses was 0.09 in the most proximal zone (0-25 nm from the AZ PM), and peaked to 0.18 in 25-50 nm zone. It then dropped to 0.08 in the intermediate zone (50-75 nm) and rose steadily more distally to reach a plateau of ~0.16 spanning the range of 150 to 250 nm distance. Finally, SV occupancy gradually decreased as the distance from the AZ increased (Figure 3A).

**Figure 3:**
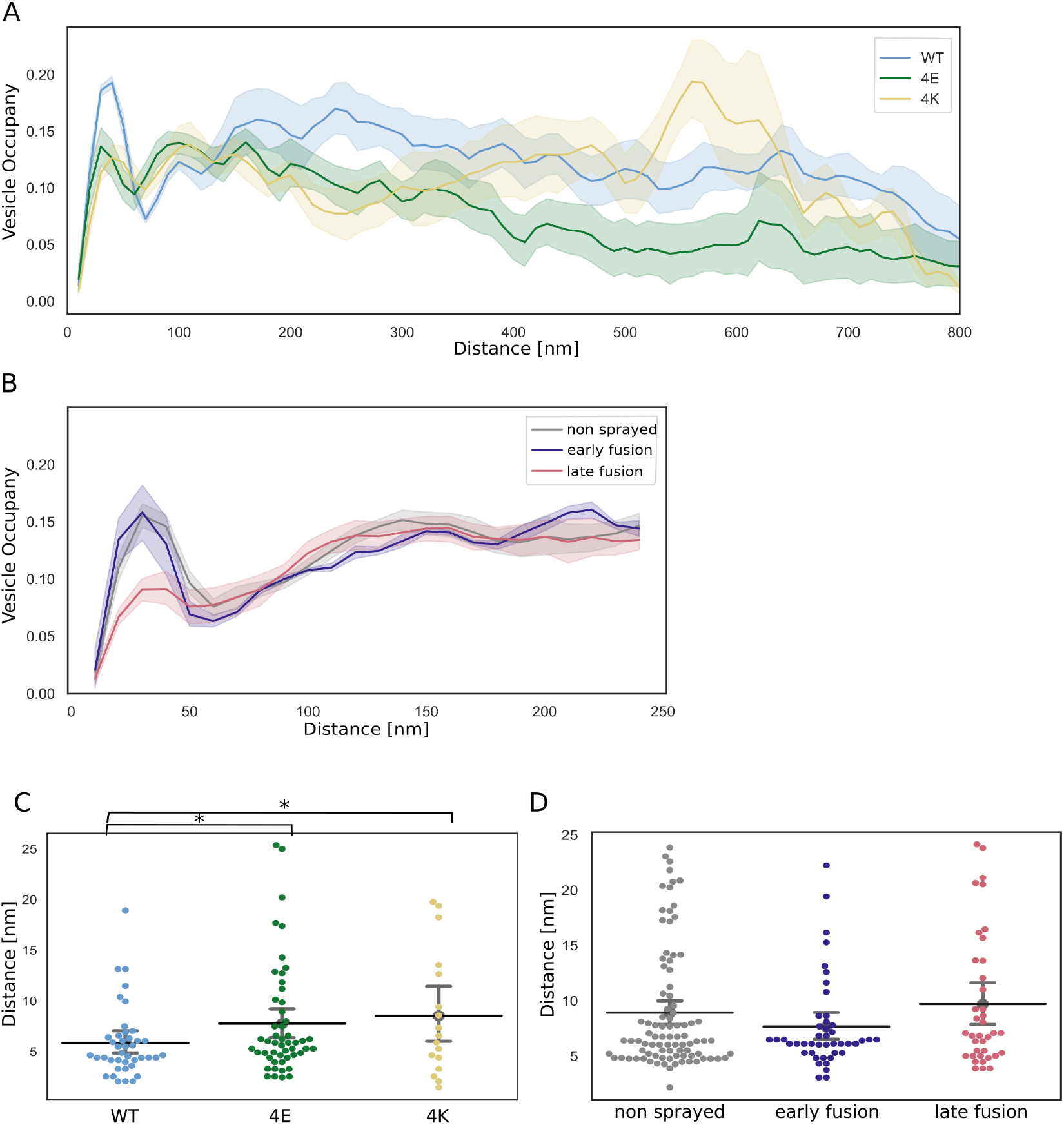
SV distribution. (A, B) Vesicle occupancy expressed as fraction of cytosol volume occupied by vesicles as a function of distance to AZ in (A) cultured neurons and (B) synaptosomes. Each solid line represents the mean occupancy value for the experimental group, while the shaded areas depict twice the standard error of the mean (SEM). (C, D) Distance of proximal SVs from the AZ. Horizontal line: mean; whiskers: 2xSEM interval. Statistical test: multiple all against control pairwise ANOVA comparisons with Benjamini-Hochberg correction; *: P<0.05

The absolute values differ between WT cultured mouse neurons and non-stimulated rat synaptosomes but the SV occupancy distribution follows the same pattern. The difference in absolute values can likely be attributed to the different experimental and animal models used. Sprayed synaptosomes that were showing early signs of exocytosis had a nearly identical SV occupancy pattern as non-sprayed synaptosomes (Figure 3B, dark blue and gray, respectively). However, when SV full collapse figures were apparent, SV occupancy in the proximal zone was significantly reduced, whereas SV occupancy further away from the AZ PM was unchanged (Figure 3B, red). This is consistent with some membrane proximal SVs having engaged in exocytosis, while none of the recycling and reserve pool SVs have. In order to investigate the consequences of chronic high or low synaptic activity, we investigated the 4E and 4K mutants (Figure 3A, green and gold, respectively). In the most proximal 50 nm, SV were significantly less concentrated in the constitutively depressed 4E mutant than in the WT. However, they were significantly more abundant between 75 and 100 nm. Furthermore, proximal SVs were on average more distant to the AZ PM in the 4E mutant than in the WT (Figure 3C). These observations are consistent with the increased distance between SVs and the PM induced by the repulsion between negative charges present in SNAP-25 and on the AZ/PM. In the most distal zones, SV occupancy gradually decreased in the 4E mutant and was significantly lower than in the WT over most of the 250 to 750 nm distance range. The decrease may reflect deleterious effects associated with abnormally low synaptic activity. The 4K mutant displayed a significantly decreased SV occupancy in comparison to the WT in the most proximal 50 nm. 4K proximal SVs were in average more located further away from the AZ PM than WT proximal SVs. Furthermore, proximal SVs were in average more distant to the AZ PM in the 4K mutant than in the WT (Figure 3C). This can be readily attributed to the high probability of spontaneous exocytosis generated by the additional positive charges of the SNARE bundle. Between 50 nm and 75 nm away from the AZ PM, SV occupancy was higher in the 4K mutant than in the WT, consistent with recycling pool SVs being recruited. From 100 to 250 nm, SV occupancy dropped steadily, in contrary to the WT, and from a distance of 170 nm, it was significantly lower. Yet, beyond 250 nm, it rose linearly until 450 nm, becoming indistinguishable from WT occupancy, and then experienced a sharp increase, peaking to 0.2 at a distance of 550 nm, significantly higher than WT occupancy, before dropping quickly.

### Proximal vesicles form additional tethers following stimulation

We investigated the tethering state of proximal SVs (i.e. the SVs whose center is located within 45 nm of the AZ PM) prior to and following stimulation in synaptosomes. In non-sprayed synaptosomes, 54% of the proximal vesicles were tethered, which is in agreement with previous results (Supplementary Figure S4B) [13].

Interestingly, in the early fusion group the fraction of tethered proximal vesicles significantly increased to 80% (P<0.05, χ2 test). In the late fusion group, however, 53% of the proximal vesicles were tethered, which is not significantly different to the non-sprayed group. The average number of tethers per proximal SV followed the same pattern. Proximal SVs had 0.89 ± 0.12 tethers in the non-sprayed group (Figure 4D). This parameter rose to 2.09 ± 0.33 in the early fusion group, while it returned to 1.00 ± 0.20 in the late fusion group.

**Figure 4:**
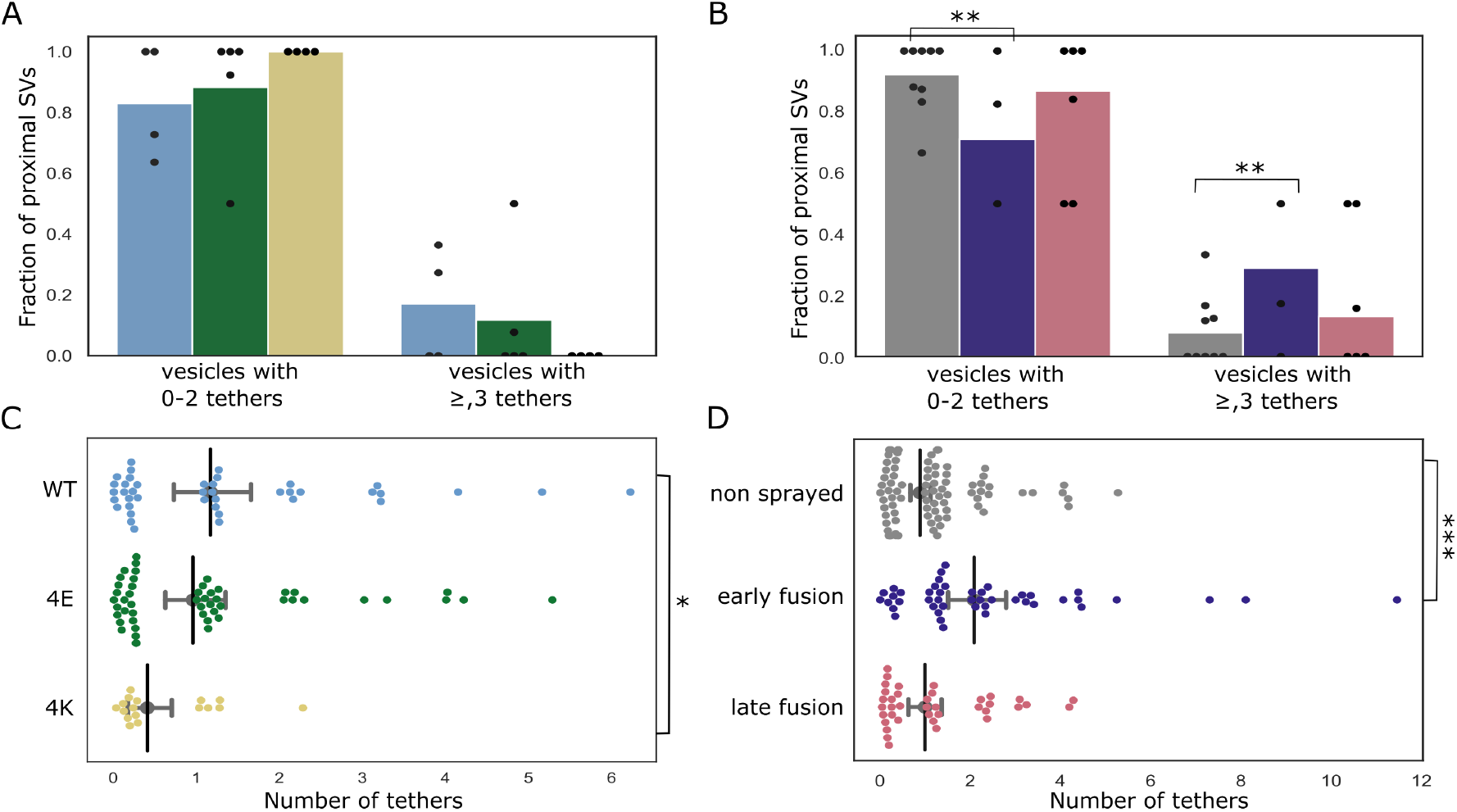
Proximal SV tethering. (A, B) Fraction of proximal SVs that are triple tethered. Each bar shows the overall fraction of all proximal SVs from a given experimental condition. Each dot represents the value of an individual synapse. Statistical test: multiple all against control pairwise χ^2^-test with Benjamini-Hochberg correction. (C, D) Number of tethers per proximal SV. Each dot represents an individual SV. The vertical line represents the mean value, and the horizontal whiskers correspond to the 95% confidence interval. Statistical test: multiple all against control pairwise ANOVA comparisons with Benjamini-Hochberg correction; *: P<0.05, **: P<0.01, ***: P<0.001.

We then analyzed whether the decreased occupancy in the late fusion group was associated with a decreased number of triple-tethered SVs, (defined as SV with at least three tethers) which as mentioned in the introduction are suggested to belong to the RRP. In resting, non-sprayed synapses 8% of the proximal SVs were triple-tethered (Figure 4B). Surprisingly, the fraction of triple-tethered proximal SVs drastically increased to 29% in the early fusion group (P<0.01, χ2 test). The fraction decreased to 13% in the late fusion group. This suggests that upon stimulation some proximal SVs very rapidly acquire new tethers. Using our definition of the RRP (vesicles that are triple-tethered) this would indicate that the RRP rapidly increases after stimulation and more vesicles become primed for exocytosis. Furthermore, the lower proximal vesicle occupancy in the late fusion group indicates that under our stimulation conditions, replenishing vesicles to the proximal zone is slower than their release.

The situation in the WT-SNAP-25 neurons was similar to unstimulated synaptosomes. 53% of the all proximal SVs were tethered and 17% of all proximal SVs belonged were triple-tethered (Supplementary Figure S4A and Figure 4A. On average, proximal SVs had 1.17 ± 0.23 tethers. The corresponding values for the 4E mutants were not significantly different (15% and 0.96 ± 0.18, respectively). However, in all 4K mutant datasets there was not a single SV that was part of the RRP, i.e. triple-tethered. Consistently, the number of tethers per proximal SV was significantly lower in the 4K mutant than in the WT (Figure 4C). These results are in line with physiological measurements that have shown that the RRP is depleted in the chronically spontaneously active 4K mutant, and they provide additional evidence that RRP-vesicles have at least 3 tethers. [12].

### Synaptic activity modifies inter-SV connectivity

The majority of SV are linked to other SVs via molecular bridges previously termed connectors [13,14]. The function and composition of connectors are not clear yet. It was earlier proposed that connectors limit SV dispersion and allow SV mobilization for release. It is generally assumed that synapsin is involved in connector formation and may be one of its components. It has been suggested that connectors reduce SV mobility and maintain a local high SV concentration in the presynapse. The connectivity level of an individual SV might be one of the factors defining the pool to which the SV belongs. To shed some light on the role of connectors, we analyzed SV connectivity in our datasets. We focused most of our analysis to the SVs located at distance of the AZ PM lower than 250 nm. Furthermore, we defined 4 distance groups: proximal (0-45 nm), intermediate (45-75 nm), distal 1 (75-150 nm), distal 2 (150-250 nm), as in previous studies [13,32]. We first analyzed synaptosomes. In non-sprayed synaptosomes datasets, approximately 70% of the proximal and intermediate SVs were connected to other vesicles. In distal 1 and 2 regions, this value rose to 84% (P<0.05) and 87% (P<0.01), respectively; Figure 5D). Similarly, the number of connectors per vesicle significantly increased from the proximal region (1.63 ± 0.13) to the distal 1 region (2.57 ± 0.09, P<0.0001) and the distal 2 region (2.78 ± 0.10, P<0.0001) (Figure 5B). Sprayed early and late fusion synapses showed a similar pattern, with significantly more connectors per SV in distal 1 region than in proximal region (P<0.01 and P<0.0001, respectively), and more connectors per SV in distal 2 region than in the proximal region (P<0.01 and P<0.0001, respectively). We then compared the number of connectors per SV between non-sprayed synaptosomes and early fusion or late fusion synaptosomes. In the proximal group no significant difference between the control and the early fusion group (1.63 ± 0.13 and 2.24 ± 0.33, P = 0.13) but it significantly dropped to 1.12 ± 0.15 in the late fusion group (P<0.05, Figure 5B). Consistently, the number of connectors per non triple-tethered proximal SV went from 1.64 ± 0.17 in the non-sprayed group, rose significantly to 2.69 ± 0.54 in the early fusion group (P<0.05) and dropped to 0.9±0.19 in the late fusion group (P<0.05) (Figure 5F). Taken together, our observations indicate that following depolarization, the number of connectors per proximal SV with less than 3 tethers (i.e. non-RRP) first increases and then decreases to a value lower than the initial one. We have seen earlier that the fraction of tethered proximal SVs does not differ between non-sprayed and late fusion synaptosomes. Thus, our data suggest that establishing connectivity is a slower process than tethering. We hypothesize that given the free space made in the proximal region after some SVs have fused, non-connected vesicles from the intermediate region diffuse to the proximal zone and become tethered to the AZ PM. Only subsequently, these newly tethered vesicles get interconnected. Furthermore, we have observed that connectors remained present between fusing SV and neighbor SV (Supplementary Figure S5). This, in addition to passive diffusion, can contribute to replenishing the RRP.

**Figure 5:**
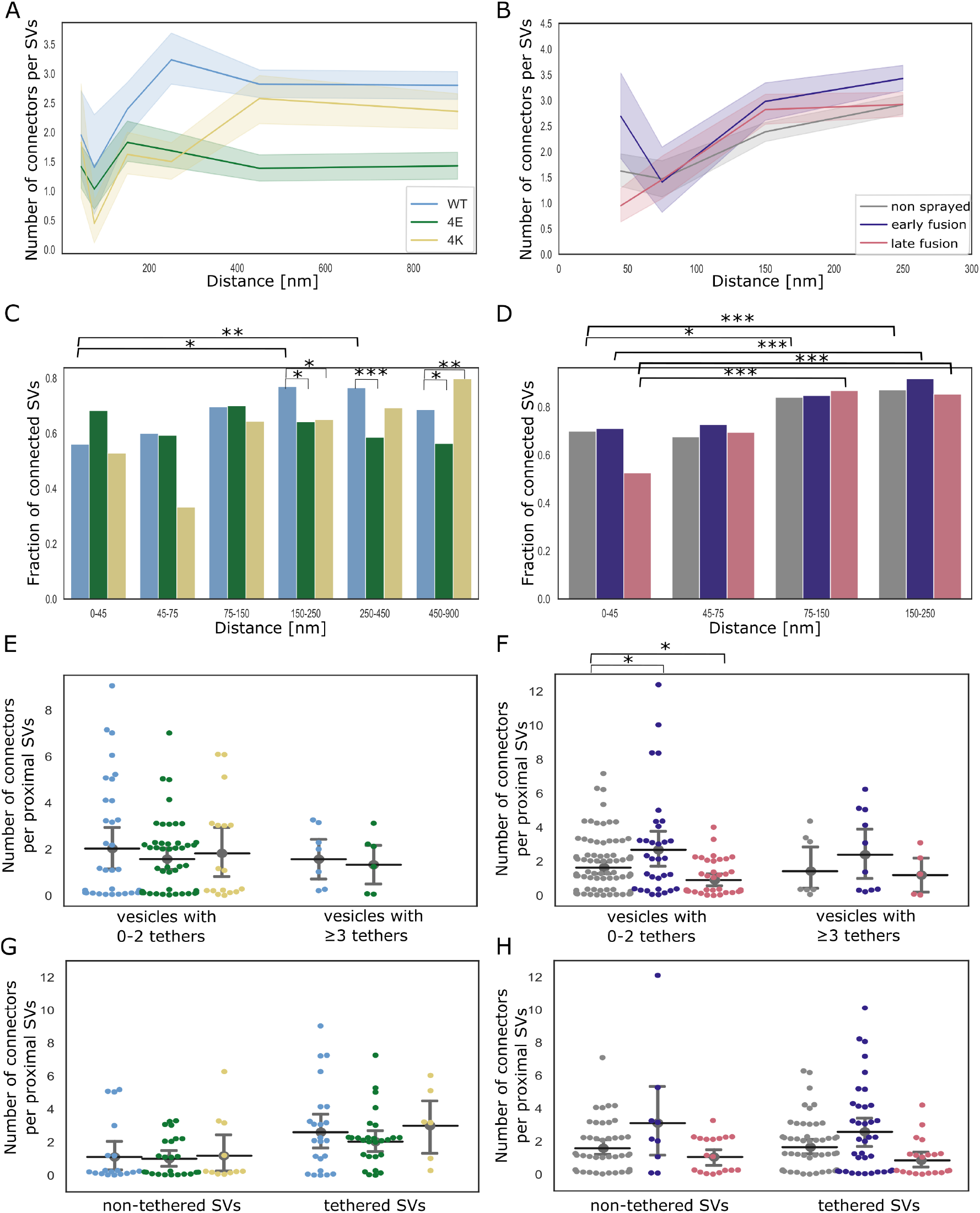
SV connectivity. (A, B) Number of connectors per SV as a function of their distance to the AZ PM for mouse neurons (A) and rat synaptosomes (B) Each solid line represents the average value of all SVs belonging to a particular experimental condition. Shaded areas represent 2xSEM interval ranges. Statistical tests: multiple all against reference pairwise ANOVA comparisons with Benjamini-Hochberg correction. Within a single experimental condition, the reference was the proximal distance group; within a single distance group, the reference was the WT genotype or non sprayed synaptosomes. (C, D) Fraction of connected vesicles as a function of distance to the AZ PM for mouse neurons (C) and rat synaptosomes (D). Statistical test: multiple all against reference pairwise χ^2^-test with Benjamini-Hochberg correction; references were defined as in (A) and (B). (E, F) Number of connectors per proximal SV belonging or not to the RRP for mouse neurons (E) and rat synaptosomes (F). (G, H) Number of connectors per tethered or non-tethered proximal SV for mouse neurons (G) and rat synaptosomes (H). Statistical tests in (E-H): multiple all against control pairwise ANOVA comparisons with Benjamini-Hochberg correction. Control was WT genotype or non sprayed synaptosomes. In all statistical tests, *: P<0.05, **: P<0.01, ***: P<0.001.

We then analyzed SNAP-25 neurons. For SNAP-25-WT, similarly to non-sprayed synaptosome, the fraction of connected SVs was significantly higher in the distal 2 region than in the proximal region (p<0.01), albeit the absolute values were overall lower than in synaptosomes (Figure 5C). Consistently, the number of connectors per SV in SNAP-25-WT synapses increased from 1.95 ± 0.38 in the proximal region to 3.23 ± 0.21 in the distal 2 region (Figure 5A, P<0.01). The fraction of connected SVs in the distal 2 region was significantly lower in the 4E and 4K mutant than in the WT (p<0.05). This was supported by a significantly lower number of connectors per SV in the distal 2 region for the 4E mutant versus the WT (P<0.0001) as well as for the 4K mutant versus the WT (P<0.0001, Figure 5A). The number of connectors per proximal SV was not affected by the mutations (Figure 5E and G). These results indicate that prolonged abnormal exocytotic activity is correlated with severe changes in intervesicular connectivity in the distal region.

## Discussion

Due to its transient nature, SV exocytosis has been difficult to characterize morphologically. A number of questions remain partially unresolved to this date. In particular, it has been suggested that following Ca^2+^ entry, the insertion of synaptotagmin-1 into the membrane induces an increase in membrane curvature, which lowers the energy barrier of fusion. Such membrane deformations have been observed in biochemically reconstituted models of exocytosis but have not yet been reported in functional synapses [11,33]. Moreover, it is not clear whether the membrane deformation occurs subsequently to Ca^2+^ influx or if primed SVs and their PM counterpart present such deformation [33]. The optimal sample preservation delivered by cryo-ET makes it possible to investigate the role of tethers located between SVs and the AZ PM and the function of inter-SV connectors. Combining cryo-ET with spray-mixing plunge-freezing enabled us to investigate the morphological changes occurring immediately after depolarization.

### Membrane curvature increases following depolarization

Depolarization through spraying droplets of KCl solution on synaptosomes milliseconds before freezing allowed us to capture snapshots of exocytosis (Figure 2B1-B3). In spite of the uncertainty on the exact delay between stimulation and freezing, our approach allowed access to shorter delays than any other time-resolved cryo-EM technique. The temporal sorting of observed exocytosis snapshots was done in the most parsimonious way. We observed that the curvature of some PM regions facing some SVs increased following depolarization. The SV facing such a PM buckling also seemed to get kinked. These deformations were not seen in non-sprayed synaptosomes. This indicates that in functional synapses exocytosis starts with a Ca^2+^-dependent membrane deformation, which is supported by a wealth of in vitro biochemical data [11,34]. Deformation may be caused in part by the intercalation of synaptotagmin-1 C2A and C2B domains between membrane head groups. A recent biophysical study indicated that C2A and C2B preferably insert in SV membrane and PM, respectively [35]. It may also be due to the tension/force induced by SNARE-complex zippering [36]. Subsequent snapshots showed a fuzzy contact point between the SV and the PM, which likely corresponds to lipid splaying or the merging of the two membranes. Membrane fusion then occurred and yielded classical Ω-fìgures with variable neck diameters. Finally, nearly fully collapsed SVs were imaged. Overall our observations support the standard model of full collapse SNARE-dependent membrane fusion [37,38] and reveal details of exocytosis early stage, prior to actual membrane fusion.

### SV local concentration correlates with SV connectivity

SV local concentration - a.k.a SV occupancy - is tightly correlated with the distance from the AZ PM. Under resting conditions, SV occupancy reaches a local maximum at the distance approximately 25 nm and a local minimum at approximately 75 nm from the AZ PM, before rising again with the distance increasing (Figure 3A and B), in agreement with previous reports [13]. By definition, all SVs in the proximal region are directly facing the PM. Their high concentration can be attributed to the fact that more than 50% of them are tethered to the PM. On the other hand, the number of connectors per SV and SV connectivity gradually increases with the distance from the AZ (Figure 5A-D). This increase correlates with the increase in occupancy. Thus, we may hypothesize that SV local concentration is a function of their level of tethering to the PM and of connection with other SVs.

Interestingly, under short stimulation of a few milliseconds, SV occupancy only decreases in the proximal region, as a consequence of the fusion of SVs with the PM (Figure 3B). In order to further assess the relation between SV tethering, connectivity, and occupancy, we analyzed synapses bearing expressing either WT SNAP-25, a more positively charged mutant (4K), or a more negatively charged mutant (4E) [12]. The 4K mutant has a decreased energy barrier to membrane fusion and causes constitutively active exocytosis, whereas the 4E mutant shows a decreased exocytotic activity because of a higher energy barrier to membrane fusion. Both mutants had a significantly decreased proximal SV occupancy (Figure 3A). In the case of the 4K mutant, this was probably due to the high frequency of spontaneous exocytosis. On the other hand, the 4E mutant, because of its additive negative charges, tends to repel SVs from the PM (Figure 3C), which can explain their decreased proximal occupancy. Over a narrow distance range, around 75 nm, SV occupancy of both mutants was significantly higher than that of the WT. Further away, SV occupancy was most significantly lower for the 4E mutants than for the WT. The 4K mutant followed the same trend but had a massive increase of SV occupancy between 500 and 600 nm, before falling again. Interestingly, the number of connector per SV follows a similar pattern as SV occupancy in both mutants. In the 4E mutant, it remains significantly lower than in the WT from approximately 200 nm and beyond. In the 4K mutant, this value is significantly lower than in the WT from approximately 100 nm but it then rises close to WT values from approximately 400 nm onward. This increased occupancy may be due to the recycling of spontaneously fusing SVs. Our data show that strong disturbances in exocytotic activity lead to profound differences in SV occupancy and SV connectivity. We note that a correlation exists between SV connectivity and concentration. Future studies will be necessary to assess whether SV concentration depends on the SV connectivity and to decipher the molecular mechanism influencing these parameters.

### SNAP-25 4K mutant further supports the RRP morphological definition

Previously, we showed that the number of tethers of a SV defines whether its exocytosis can be induced by treatment with a hyperosmotic sucrose solution, which corresponds to a definition of the RRP [13,14]. We reported that SVs with at least 3 tethers belong to the RRP, according to this definition. In order to further assess this model, we analyzed synapses of neurons expressing the SNAP-25 mutants. 17% of the WT proximal SVs had 3 tethers or more. Critically, the 4K mutant had none such SV. As the RRP (assessed with hyperosmotic sucrose treatment) in this mutant was formerly shown through functional assays to be depleted, our present observation further supports our morphological definition of the RRP [12]. 15% of the proximal SVs had 3 tethers or more in the 4E mutant, which is very similar to the WT situation, while this mutant was shown to possess a normal-sized RRP. Our observations are also consistent with a number of studies that have concluded that SV exocytosis requires a minimum of three SNARE complexes [8,9,10].

### Depolarization rapidly induces additional tethering in proximal vesicles

We compared SV tethering before and shortly after depolarization. Our observations are schematically summarized in Figure 6. Interestingly, the fraction of proximal SVs that were tethered increased by 50% shortly after stimulation, in synapses showing early signs of exocytosis. Simultaneously, the number of tethers per proximal SV more than doubled and, the fraction of proximal SVs with 3 or more tethers tripled. In presynaptic terminals presenting more advanced stages of exocytosis (Ω-figures), all these measurements returned to pre-stimulation values. These data indicate that immediately after the onset of stimulation a quick and massive increase in tethering occurs. This phenomenon was resolved in our measurements, because the spraying of synaptosomes with an intermediate K^+^-concentration made it possible to isolate synaptosomes in an early stage of fusion, which would have been missed during either strong or chronic stimulation, which would deplete primed vesicles.

**Figure 6:**
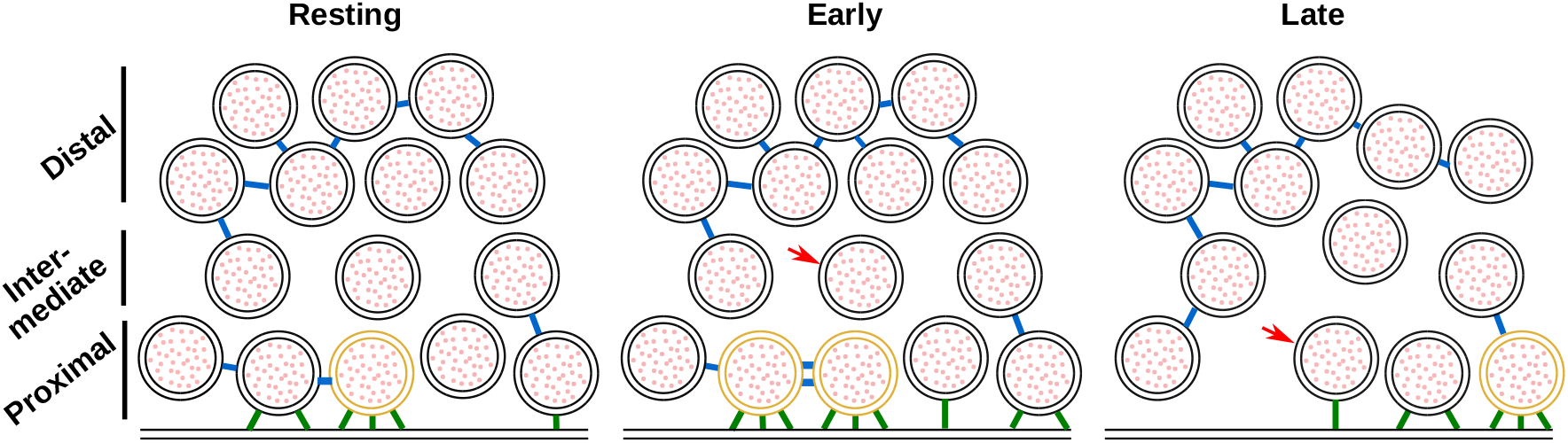
Model depicting a synapse transitioning from resting state to early and late fusion states. Tethering and connectivity changes upon synapse stimulation are depicted. Proximal non triple-tethered vesicles (black proximal SVs) gain additional tethers and some of them become triple-tethered (yellow SVs) shortly after stimulation. Primed vesicles then fuse with the plasma membrane (late fusion) and leave an empty space in the AZ cytoplasm. The number of connectors (depicted in blue) per proximal SV decreases in late fusion tripled-tethered vesicles. The red arrow shows a vesicle initially located in the intermediate region, which diffuses to the proximal region in the late fusion state. Tethers are shown in green.

The phenomenon of rapid, depolarization-induced tethering leads to some free proximal SVs becoming tethered to the AZ PM, while some previously single- or double-tethered SVs gained the additional tether(s) that according to our definition of the RRP (as triple-tethered vesicles) would be expected to render them releasable [13]. There are several important implications of this finding. First, the increase of the number of tethers during the initial membrane contact - in excess of the three tethers formed during priming - might help overcome the fusion barrier. Functional reconstruction led to the suggestion that SNARE-complexes primarily form downstream of Ca^2+^-influx [39], whereas mutagenesis studies in cells supported the notion that SNARE-complexes had already formed before arrival of the Ca^2+^-trigger, i.e. during priming [40]. In fact, both notions might be partly correct, as the formation of a low number of SNARE-complexes might lead to a stable primed state, defined by a valley in the energy landscape due to the dual inhibitory/stimulatory features of the SNARE-complex [12,41], whereas more SNARE-complexes might form dynamically after triggering, during membrane fusion itself. Accordingly, in in vitro fusion assays additional SNARE-complexes, above those required for fusion pore formation, leads to fusion pore stabilization and release of larger cargos [10,42]. Second, vesicles that have not formed three tethers before stimulation might fuse with delayed kinetics during triggering, which accounts for the variable exocytosis kinetics among SVs [43,44,45,46]. Superprimed vesicles are expected to have formed the largest number of tethers before stimulation [43,44]. Third, overlapping protein complexes might be involved in priming and triggering, depending on the timing of their formation. Accordingly, triggering that stimulates tether-formation might also stimulate priming for those vesicles that were not tethered before stimulation. Indeed, a number of recent publications have suggested that some SVs can get primed extremely quickly in response to Ca^2+^ influx [46,47,48,49,50].

### Conclusion

Our study revealed fine morphological changes occurring in the presynaptic terminal immediately after the onset of exocytosis, as well as in chronically active or inactive synapses. It indicates increased SV tethering induced the rise in presynaptic Ca^2+^, potentially corresponding to SV superpriming, and preceding SV fusion. It also highlights modifications of proximal SV interconnections in response to evoked exocytosis, as well as more drastic modifications of distal SV interconnections in chronically active synapses and in inactive synapses. These changes likely affect SV mobility and recruitment at the AZ.

## Materials and methods

### Constructs and viruses

SNAP-25B was N-terminally fused to GFP and cloned into a pLenti construct with a CMV promoter [51]. Mutations were made using the QuikChange II XL kit (Agilent). The mutations were verified by sequencing and have been published before [12]. The preparation of lentiviral particles followed standard protocols.

### Animals

Synaptosomes were prepared from adult male or female Wistar rats obtained from the central animal facilities of the Department of Biomedical Research of the University of Bern. Adult male or female Wistar rats at an age of 6-8 weeks were slightly stunned by CO_2_ and quickly decapitated with a guillotine. The procedures used were in accordance with the Swiss Veterinary Law guidelines. Heterozygous SNAP-25 KO C57/Bl6-mice were routinely backcrossed to Bl6 to generate new heterozygotes. The strain was kept in the heterozygous condition and timed heterozygous crosses and caesarean section were used to recover knockout embryos at embryonic day 18 (E18). Pregnant females were killed by cervical dislocation; embryos of either sex were collected and killed by decapitation. Permission to keep and breed SNAP-25 mice was obtained from the Danish Animal Experiments Inspectorate and followed institutional guidelines as overseen by the Institutional Animal Care and Use Committee (IACUC). Newborn (P0-P2) CD1 outbred mice of either sex were used to create astrocytic cultures and for that were killed by decapitation.

### Synaptosome preparation

Rat synaptosomes were prepared as previously described [52], with some modifications. The cerebral cortex and the hippocampi were removed in sucrose buffer (SEH: 0.32 M sucrose, 1 mM EDTA, 10 mM HEPES; HEPES, #H4034, Sigma-Aldrich Corporate Offices. St. Louis, MO, USA) on ice. Homogenization of the tissue was done in SEH with a Potter-Elvehjem grinder (#358011, Wheaton. Millville, New Jersey, USA), four strokes at the bottom and 6 from top to bottom were applied to the tissue at a speed of 800 turns/min as described in [52]. The whole process from decapitation to homogenization was done within 2-3 min, to obtain functional synaptosomes. Homogenized tissue was then centrifuged at 1000 g for 10 min at 4°C to remove meninges and blood vessels. The resulting supernatant containing synaptosomes, but also gliosomes and mitochondria was then added to a discontinuous, isoosmotic Percoll (#P1644, Sigma) gradient with 5%, 10% and 23% in 0.32 M sucrose, 1 mM EDTA in centrifuge tubes (#344060, Beckman Coulter). The samples were spun in an ultracentrifuge (rotor: SW 40 Ti; Beckman Coulter. Nyon, Switzerland) at 16400 rpm for 12 min at 4°C. The layer with the highest amount of functional synaptosomes was between 10-23 % [52]. The layer was carefully taken out and diluted 1:10 in HEPES buffered medium (HBM; 140 mM NaCl, 5 mM KCl, 5 mM NaHCO_3_, 1.2 mM Na_2_HPO_4_, 1 mM MgCl_2_, 10 mM Glucose, 20 mM HEPES). The obtained solution was further spun with an ultracentrifuge (rotor 45 Ti; Beckman Coulter) at 11200 rpm for 20 min at 4°C. The pellet was carefully and quickly aspirated with a Pasteur pipette to avoid mixture with the solution and then diluted in HBM.

### Preparation of astrocytic and neuronal culture

The procedure has been published before [53]. Glial cells were ready to be used after 10 days. Once they were triturated and counted with a Buerker chamber, 100,000 cells/ml were plated onto untreated 12-well plates containing DMEM supplemented with 10% foetal bovine serum (FBS), 10000 IU penicillin, 10 mg streptomycin and 1% MEM non-essential amino acids (DMEM+10% FBS). Astrocytes were isolated from CD1 outbred mice (P0-P2). Pups were killed by decapitation and heads were placed in HBSS-HEPES medium (HBSS supplemented with 1 M HEPES). The cortices were isolated from the brains and the meninges were removed (dura, pia and arachnoid mater). The cortices were chopped into smaller fragments and transferred to a tube containing 0.25% trypsin dissolved in Dulbecco’s MEM (DMEM). Fragments were incubated for 15 min at 37°C. Subsequently, inactivation medium (12.5 mg albumin + 12.5 mg trypsin-inhibitor in DMEM+10% FBS) was added and the tissue washed with HBSS-HEPES. Tissue was triturated until a smooth cloudy suspension appeared. Cells were plated in 75 cm^2^ flasks with pre-warmed DMEM, one hemisphere per flask, and stored at 37°C with 5% CO_2_. Glial cells were ready to be used after 10 days. Glial cells were washed with pre-warmed HBSS-HEPES. Trypsin was added and the flasks were incubated at 37°C for 10 min. Cells were triturated and counted with a Buerker chamber before plating 100,000 cells/ml on untreated 12-well plates containing DMEM+10% FBS. After 2 days, neurons were plated.

Hippocampal neurons were isolated from E18 SNAP-25 KO of either sex. The SNAP-25 KO pups were obtained by pairing two heterozygote animals, and the embryos were recovered at E18 by caesarean section. Pups were selected based on the absence of motion after tactile stimulation and bloated neck [54]; the genotype was confirmed by PCR in all cases. The pups were killed by decapitation and heads were put in HBSS-HEPES medium. The cortices were isolated from the brains and the meninges were removed. The hippocampi were cut from the cortices before being transferred to a tube containing 0.25% trypsin dissolved in HBSS-HEPES solution. Fragments were incubated for 20 min at 37°C. Afterwards, the tissue was washed with HBSS-HEPES. The hippocampi were triturated and the cell count was determined with a Buerker chamber. 20 μl of solution containing 250,000 cells/ml were plated onto the flame sterilized gold R2/2 or R2/1 EM grids as previously described in [53]. Following a 30-min incubation at 37°C, the grid was transferred into the 12-well plate containing the astrocytes and medium was replaced with NB medium (Neurobasal with 2% B-27, 1 M HEPES, 0.26% lutamax, 14.3 mM β-mercaptoethanol, 10000 IU penicillin, 10 mg streptomycin) for the E18 pups. Between 4 h and 1 day later, lentiviral particles carrying either SNAP-25-WT, SNAP-25-4E, or SNAP-25-4K constructs were added to the culture [12]. The cultures were incubated for 12 to 14 days before being plunge frozen.

### Plunge freezing and spray-mixing

Rat synaptosomes were prepared for plunge freezing and spray-mixing as follows. The following steps from incubation to plunge freezing were all done at room temperature (RT), equivalent to 23-25°C. The synaptosomal solution was incubated with calcein blue AM (#C1429, Molecular Probes-Thermo Fisher Scientific. Waltham, MA, USA) 30 min prior to plunge freezing to visualize the cytosol of functional – esterase containing – cellular compartments such as synaptosomes. Additionally, 1.3 mM CaCl_2_ and 10 nm gold fìducials were added (gold fiducials, #s10110/8. AURION Immuno Gold Reagents & Accessories. Wageningen, The Netherlands). CaCl_2_ is necessary to trigger exocytosis and gold fìducials are important to align the acquired tilt series for tomogram reconstruction. The sprayed solution contained 1 mM CaCl_2_ and 52 mM KCl in HBM to depolarize synaptosomes and trigger exocytosis. It also contained fluorescein (#46955, Sigma) to trace the spray droplets on the EM grid in cryo-FM. The synaptosomal solution was applied to a 200-mesh lacey finder carbon film grid (#AGS166-H2. Agar Scientific. Elektron Technology UK Ltd. Stansted, UK). Excess liquid on the grid was removed by blotting with a fìlter paper and the grid was immediately plunge frozen in liquid ethane with a homebuilt plunge freezer and was sprayed on the fly. The plunge freezer and the spraying device (atomizer) were computer controlled with a LabView script (National Instruments Corporation. Mopac Expwy Austin, TX, USA). The spraying device was set similarly to the device in [30]. Nitrogen gas pressure necessary to drive spraying was set to 2.5 bar. The grid was set to pass in front of the spray nozzle at a distance of 3-4 mm. The plunge freezer was accelerated to 0.75 m/s and the minimum spray delay was ~7 ms. The atomizer sprays scattered droplets of various size on the EM grid. During the time lapse between spraying and freezing the content of the droplets spreads by diffusion. KCl diffuses approximately 4x faster than fluorescein. Cryo-ET imaging was done within the diffusion distance of KCl but outside of the visible spray droplet because the center of the spray droplet would usually be too thick for imaging. This reduces the effective stimulation duration to anything between 0 ms and less than the given spray-freeze delay. Moreover, through diffusion, KCl concentration rapidly rises and then decreases. Hence synaptosomes are not permanently depolarized.

After 12 to 14 days of incubation grids with mouse neurons were plunge frozen with a Vitrobot (Thermofisher Scientific, Mark IV) with a blot time of 3 s and a blot force of −10. Wait time and drain time were not used. Humidity was set to 100% at 4°C. 4 μl undiluted 10 nm BSA gold tracer (Aurion) was added directly onto the grid prior to plunge freezing.

### Cryo-fluorescence microscopy

After plunge freezing, rat synaptsome samples were imaged at the fluorescent microscope under cryo conditions, with a Zeiss Axio Scope.A1, equipped with an AxioCam MRm camera (Carl Zeiss AG, Germany), and a fluorescence lamp (HXP 120 C). The correlative microscopy stage (#CMS196, Linkam Scientific Instruments, UK) was cooled down to −190°C by liquid nitrogen and the frozen EM grid was placed into the chamber of the cryostage on a bridge that was partially submerged in liquid nitrogen and was close to the objective, where the temperature was around −150°C. The filter set used for imaging fluorescein was #38 (#000000-1031-346, Zeiss) (BP 470/40, FT 495, BP 525/50; corresponds to GFP) and the one for calcein blue AM was #49 (#488049-9901-000, Zeiss) (G 365, FT 395, BP 445/50; corresponds to DAPI). The objective used was either a 10x (#420941-9911, NA = 0.25 Ph1, Zeiss) or a 50x (#422472-9900, NA = 0.55 Dic, Zeiss), the acquisition software used was AxioVision (AxioVs40Ö64 V 4.8.3.0, Zeiss) and the processing software was ZEN lite (Zeiss).

### Cryo-electron microscopy

Following cryo-FM, the rat synaptosome grids were mounted in a cryo-holder (Gatan, Pleasonton, CA, USA) and transferred to a Tecnai F20 (FEI, Eindhoven, The Netherlands) which was set to low dose conditions, operated at 200 kV, and equipped with a field emission gun. Images were recorded with a 2k x 2k CCD camera (Gatan) mounted after a GIF Tridiem post-column filter (Gatan) operated in zero-loss mode. The sample was kept at about −180°C. Tilt series were acquired using SerialEM [55] for automated acquisition recorded typically from −50° to 50° with a 2° angular increment and an unbinned pixel size of 0.75 or 1.2 nm. Due to sample thickness (400-700 nm), tomograms were usually not recorded with higher tilt angles. Defocus was set between −8 to −12 μm and the total electron dose used was about 80-100 e^-^/Å^2^. Some tomograms were acquired at a Titan Krios equipped with a K2 direct electron detector (Gatan) without energy filter. The K2 camera was operated in superresolution counting mode and between 8-40 frames per tilt angle were taken. Tilt series were acquired using the Latitude software (Gatan) for automated acquisition recorded typically from −60° to 60° with a 2° angular increment and an unbinned pixel size of 0.6 nm. Defocus was set between −8 to −12 μm and the total electron dose used was about 80-100 e^-^/Å^2^. Prior to image processing the frames at each tilt angle, frames were aligned and averaged in 2dx MC_Automator [56] with motioncor [57]. 3D reconstruction was done in IMOD [58]. The alignments were done using the automated fiducial tracking function and the 3D reconstructions were done using the weighted back projection followed by a nonlinear anisotropic diffusion (NAD) filtering. Following tomogram reconstruction only synaptosomes that fulfilled the following criteria were used: 1) even and non-broken PM, 2) synaptic cleft still attached to the presynapse, 3) spherical vesicles, and 4) a mitochondrion in the presynapse necessary to cover the energy demands of the synapse. These criteria indicate that the synaptosome is functional [59].

Cultured mouse neurons tilt series were acquired at a Titan Krios, equipped with a Falcon 3 direct electron detector (Thermofisher Scientific) without energy filter. The Falcon camera was operated in linear mode. Tilt series were acquired using the TEM Tomography software (TFS) for automated acquisition recorded typically from −60° to 60° with a 2° angular increment and an unbinned pixel size of 0.37 nm. Defocus was set between −6 to −10 μm and the total electron dose used was about 80-100 e^-^/Å^2^. Tomogram reconstruction was done as for synaptosome datasets.

### Manual and automatic segmentation procedures

Manual segmentation of SVs, mitochondria, and the active zone PM was done in IMOD (Supplementary Figure S2B and D, Supplementary Movies S1 and S2). The boundary marked the region to be analyzed by Pyto [60]. The analysis by Pyto was essentially the same as described previously [13] [60]. In short, the segmented area is divided in 1 voxel thick layers parallel to the active zone for distance calculations. A hierarchical connectivity segmentation detects densities interconnecting vesicles (so-called connectors) and densities connecting vesicles to the active zone PM (so-called tethers) (Supplementary Figure S2B and D, Supplementary Movies S1 & S2). Distance calculations are done with the center of the vesicle. Mainly default settings were used. The segmentation procedure is conservative and tends to miss some tethers and connectors because of noise. Consequently, the numbers of tethers and connectors should not be considered as absolute values but rather to compare experimental groups. All tomograms analyzed by Pyto were obtained on the same microscope with the same tilt range. The margin of error for false negatives and positives was found to be less than 10% by comparison with ground truth [60]. As it was done before, an upper limit was set between 2100 and 3200 nm^3^ on segment volume. The tomograms that were used for this analysis were binned by a factor of 2 to 3, resulting in voxel sizes between 2.1 and 2.4 nm. Tether and connector length were calculated using the midpoint method [60]. From the stimulated synaptosomes only those that showed visible signs of exocytosis were used for analysis in Pyto.

### Data analysis

If not stated otherwise data in the text are described as mean ± standard error of the mean (SEM). Wherever possible, data were presented as box plots with the following settings: orange bar, median; box extremities, lower and upper quartiles; whiskers extend up to 1.5 x interquartile range; dots, outliers. Multiple pairwise ANOVA comparisons with with Benjamini-Hochberg correction was used [61]. All against control and/or all against reference pairwise comparisons were performed. See figure legends for details. In addition, for data that required to be split into discrete bins (e.g. fraction of connected vesicles by distance to active zone), pairwaise χ2 tests with Benjamini-Hochberg correction were used. We did not apply statistical methods to predetermine sample size but similar sample sizes as previously reported have been used [13]. It was not necessary to apply randomization.

### Manuscript preparation

The manuscript was written with the open and collaborative scientific writing package Manubot [62]. The source code and data for this manuscript are available at https://github.com/aseedb/synaptic_tomo_ms.

## Supplementary Material

### Supplementary Figures

**Figure S1:**
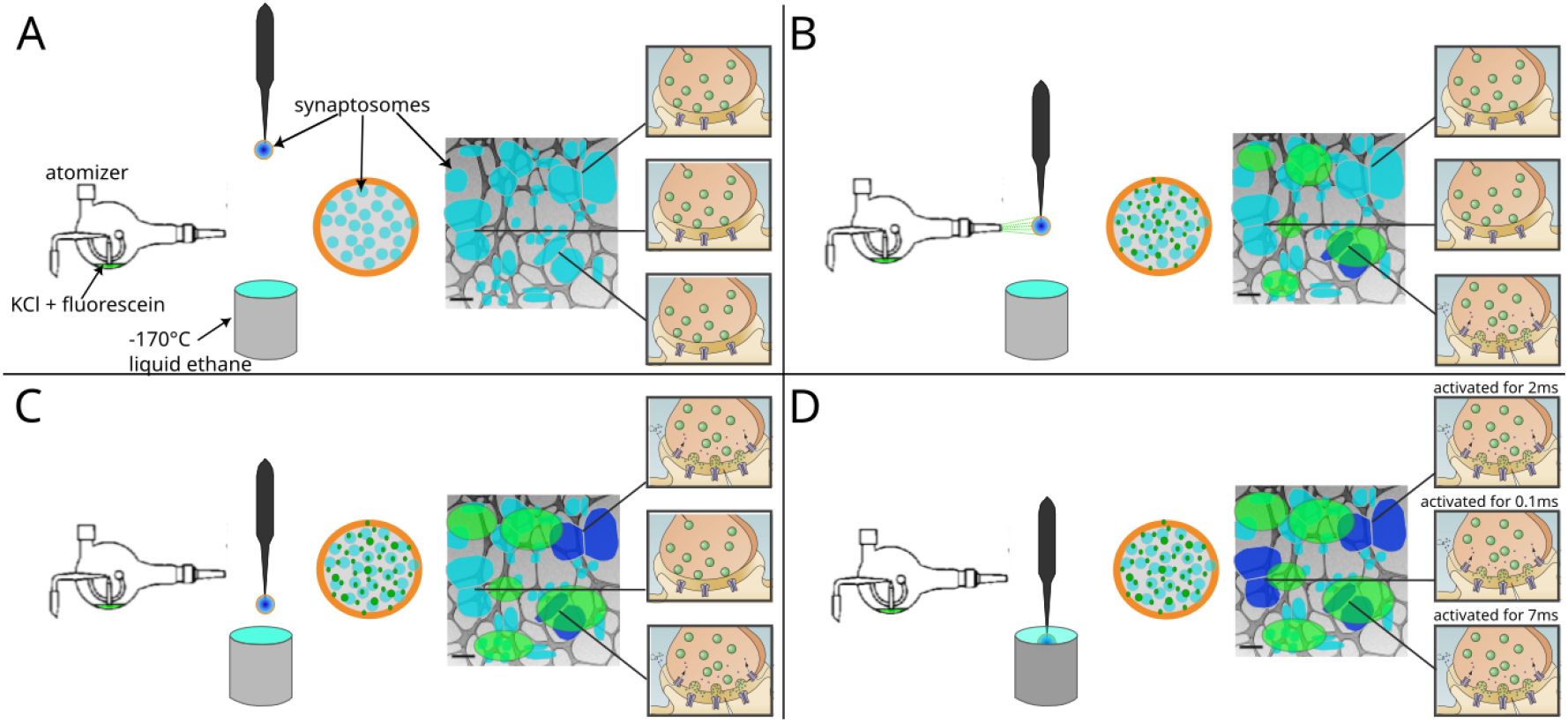
Schematic representation of a spray-mixing plunge-freezing experiment. In a single experiment different synaptosomes get stimulated for between less than 1 ms and 7 ms. An EM-grid is held by tweezers and is covered with synaptosomes in HBM-solution. A magnified view of a grid squares shows synaptosomes in blue and their synpatic state of three synapsomes is represented on the rightmost part of each panel. Panel (A) represents the situation right after blotting off solution excess. The grid is accelerated towards the spray and the cryogen. Panel (B) shows a snapshot of the experiment when the grid crosses the spray, 7 ms before the freezing. Some fluorescently dyed droplets containing HBM with 52 mM KCl land on the grid and are depicted in green. At this time point, a synaptosome located at the impact point of a droplet is activated and is depicted in dark blue. Panel (C) shows a snapshot 5 ms later, i.e. 2 ms before freezing. As KCl diffuses away from droplet impacts points, another synaptosomes gets activated because, locally KCl concentration has reached reached a concentration to depolarize enough the synaptosome so that voltage-gated calcium chanels open. Panel (D) shows a synaptosome at the time of impact with ethane. 0.1 ms before freezing a third synaptosome got exposed to a high enough concentration of KCl and got stimulated.

**Figure S2:**
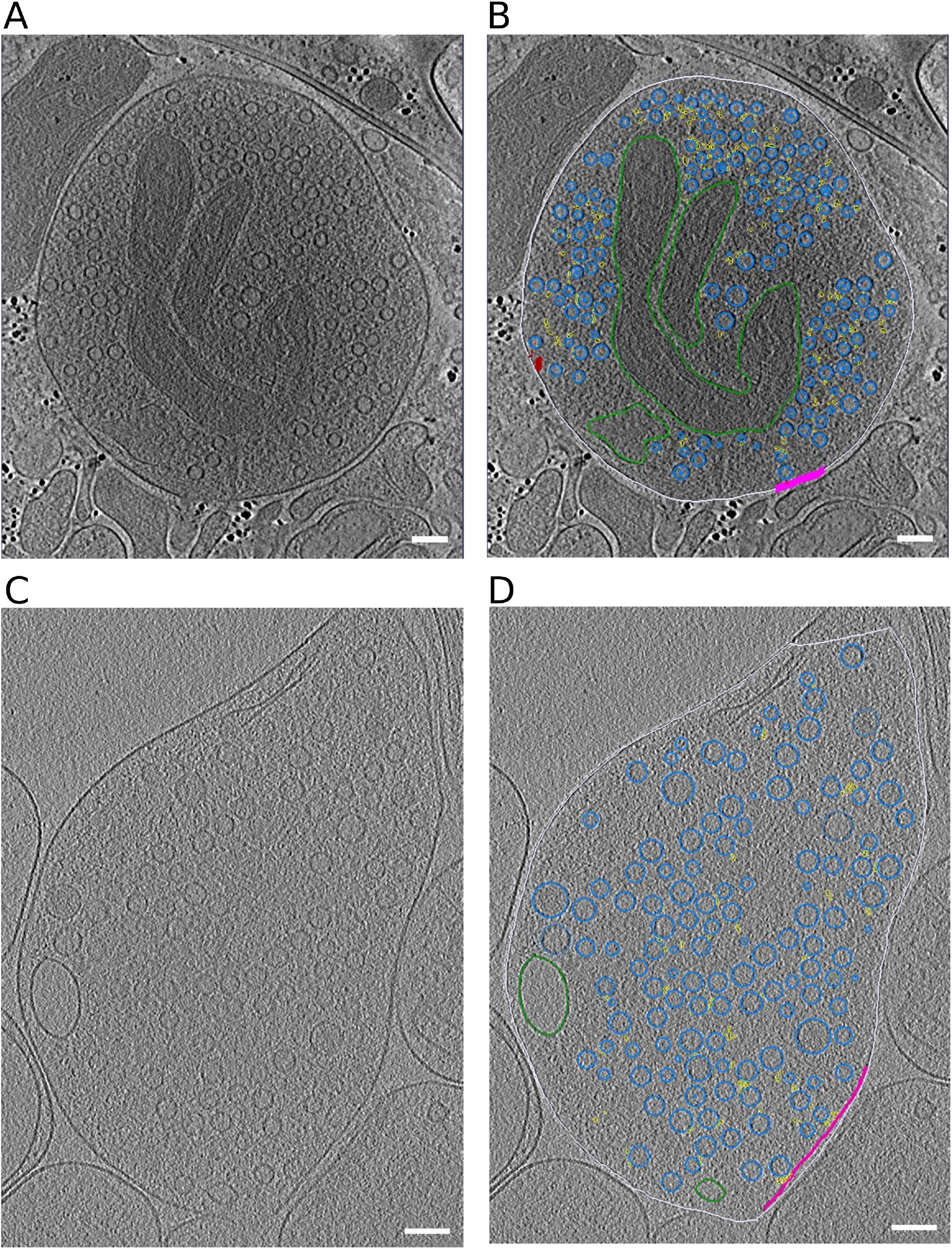
Representative slices through tomograms. (A, B) Tomographic slice without (A) and with (B) segmentation of synaptosome with late fusion events. (C,D) Tomographic slice without (C) and with (D) segmentation of WT SNAP-25 neurons. Segmentation colors: off-white = cell outline; pink = active zone; blue = synaptic vesicles; green = mitochondria; yellow = connectors, red = tethers. Scale bar, 100 nm.

**Figure S3:**
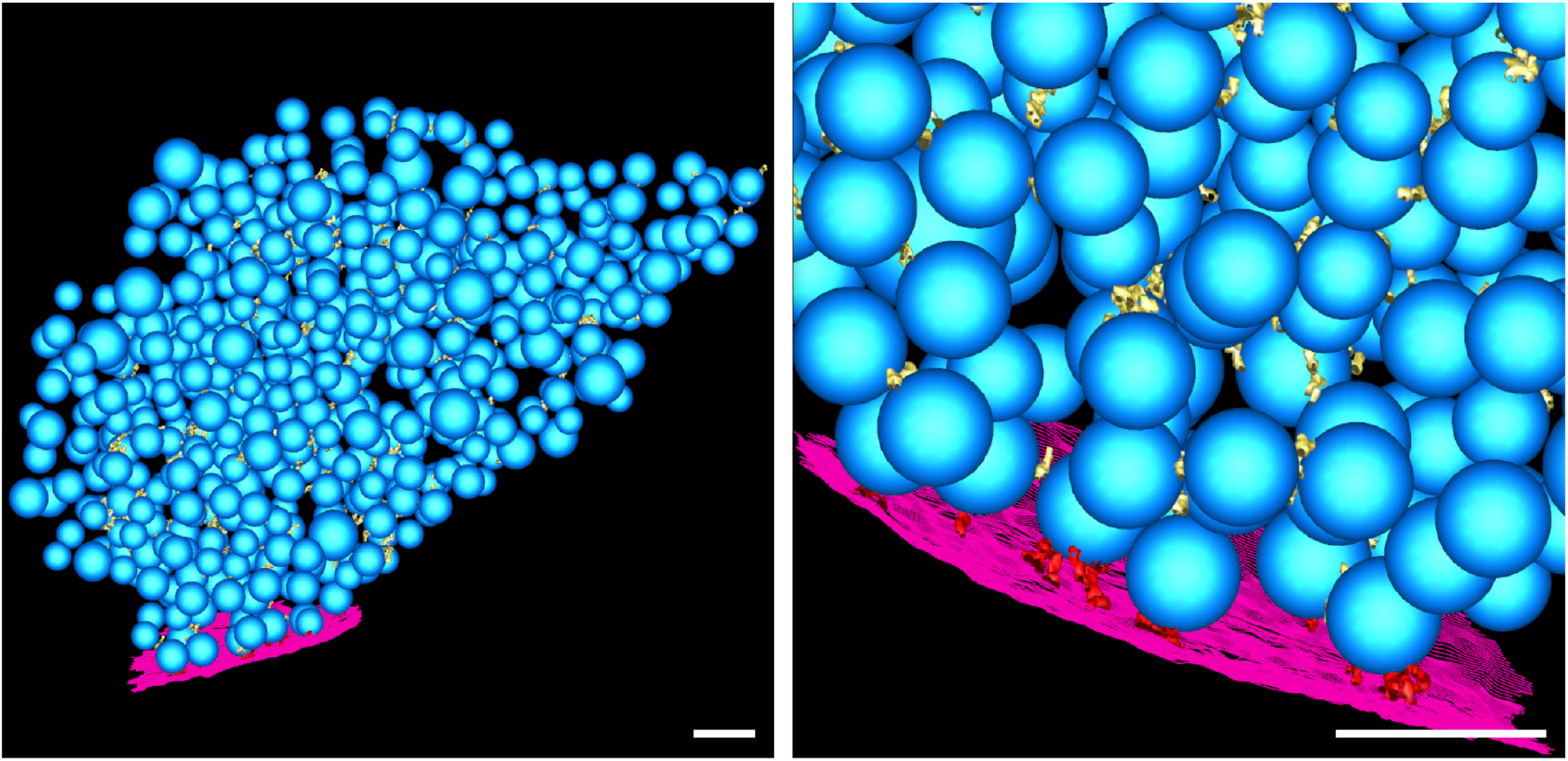
Segmented tomogram of a SNAP-25 4E neuron. (left) Overview. (right) Detail. Blue: synaptic vesicles; purple: active zone plasma memmbrane; yellow: connectors; red: tethers. Scale bar: 100 nm.

**Figure S4:**
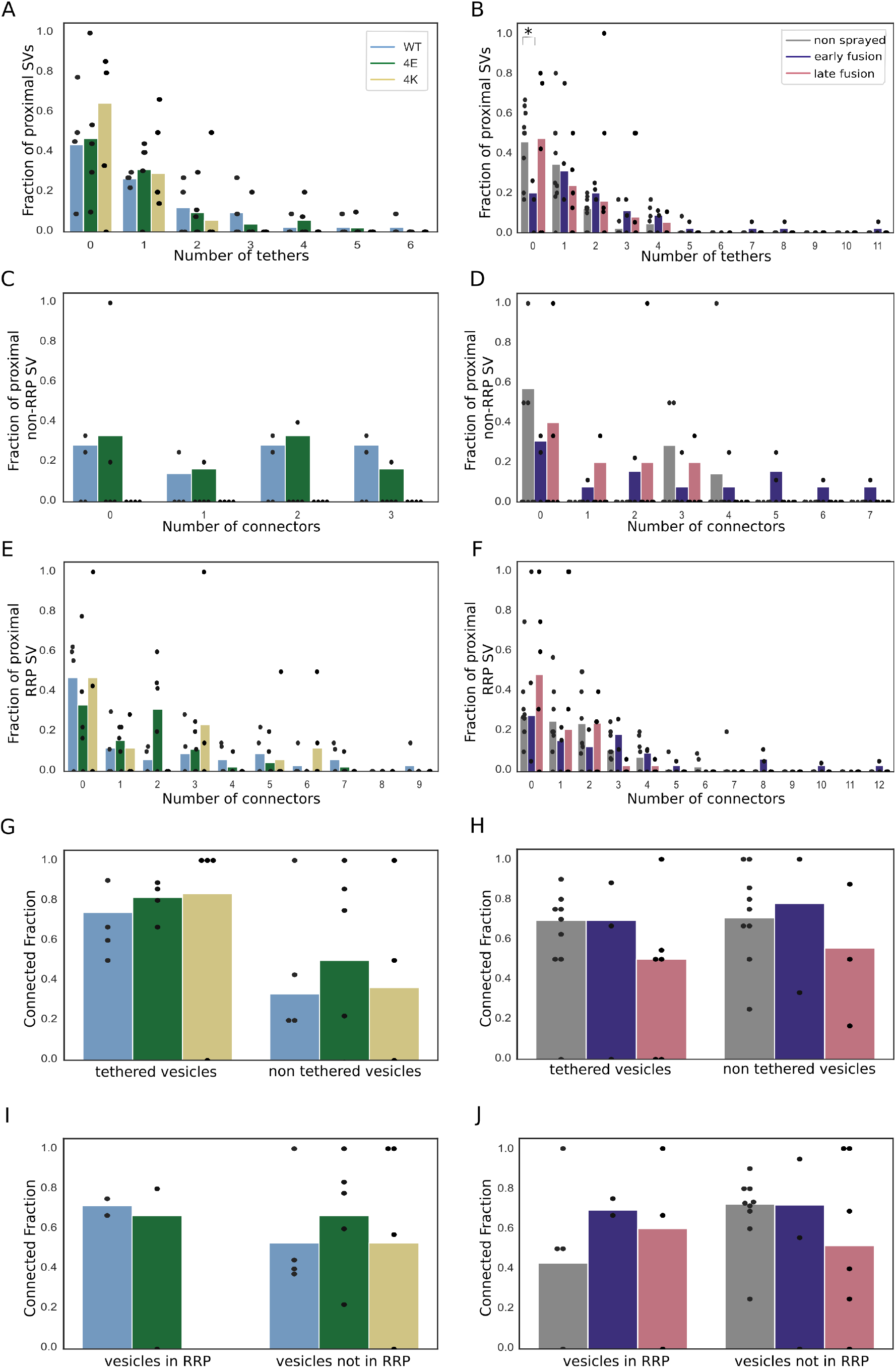
(A, B) Histogram of the number of tethers per proximal SV. Statistical test: pairwise χ^2^-test between control and each experimental condition in the 0-tether group with Benjamini-Hochberg correction. *: P<0.05. (C, D) Histogram of the number of connectors per proximal non-RRP SV. (E, F) Histogram of the number of connectors per RRP SV. (G, H) Histogram of connected SV amongst tethered or non-tethered proximal SVs. (I, J) Histogram of connected SV amongst proximal non-RRP or RRP SVs. (A, C, E, G, I) Synapses in mouse cultured neurons. (B, D, F, H, J) Rat synaptosomes.

**Figure S5:**
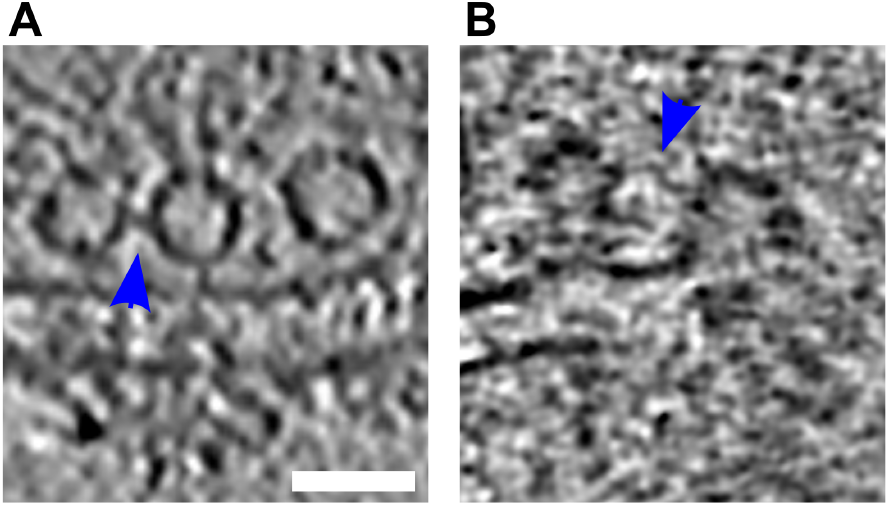
(A, B) **Tomographic slices showing tethered connected vesicles**. Blue arrows highlight the connectors. Scale bar, 50 nm.

### Supplementary Movies

**Movie S1:**
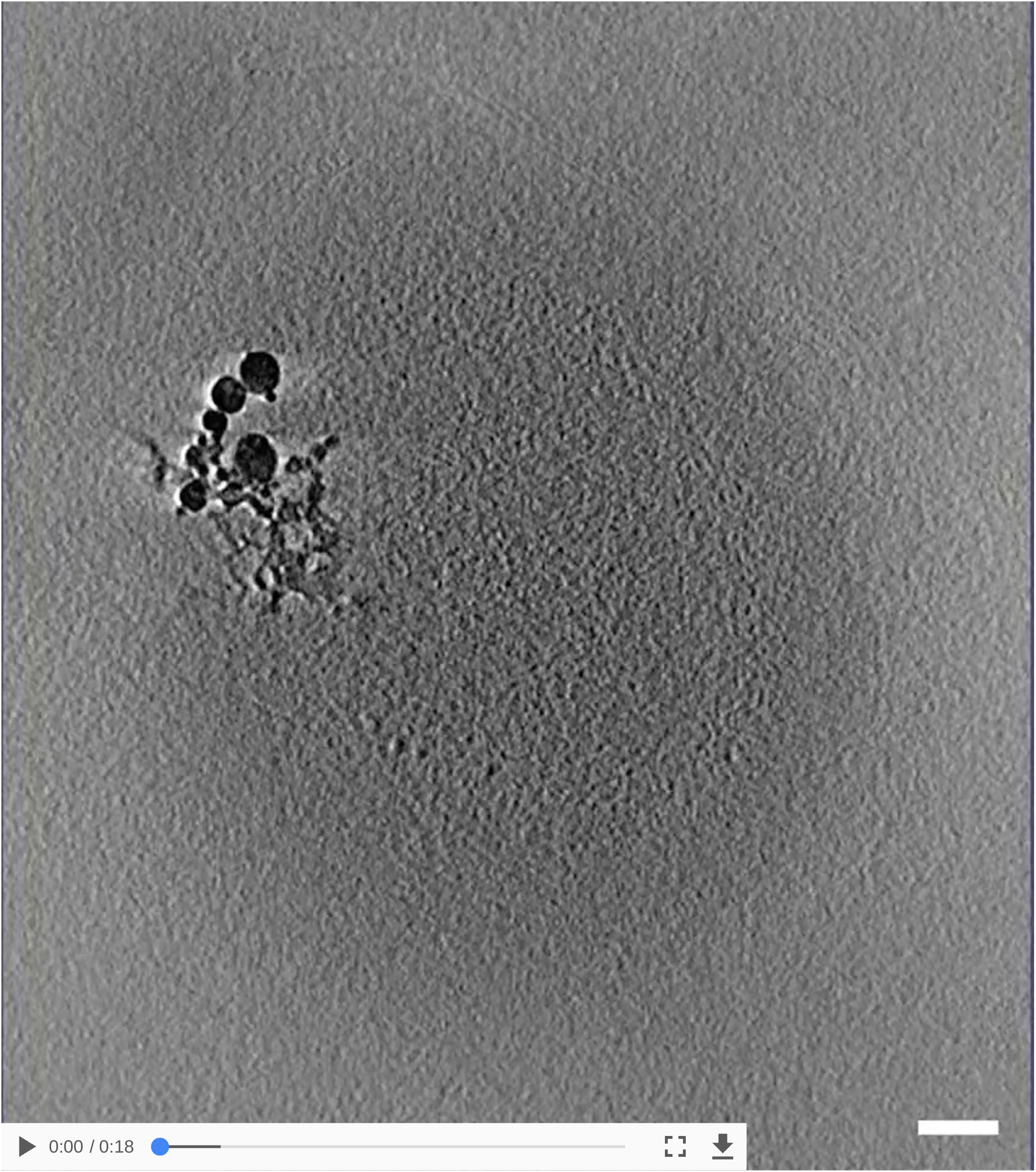
Tomogram with segmentation of synaptosome with late fusion events. off-white = cell outline; pink = active zone; blue = synaptic vesicles; dark green = mitochondria; light green = large vesicles; yellow = connectors, red = tethers, scale bar 100nm

**Movie S2:**
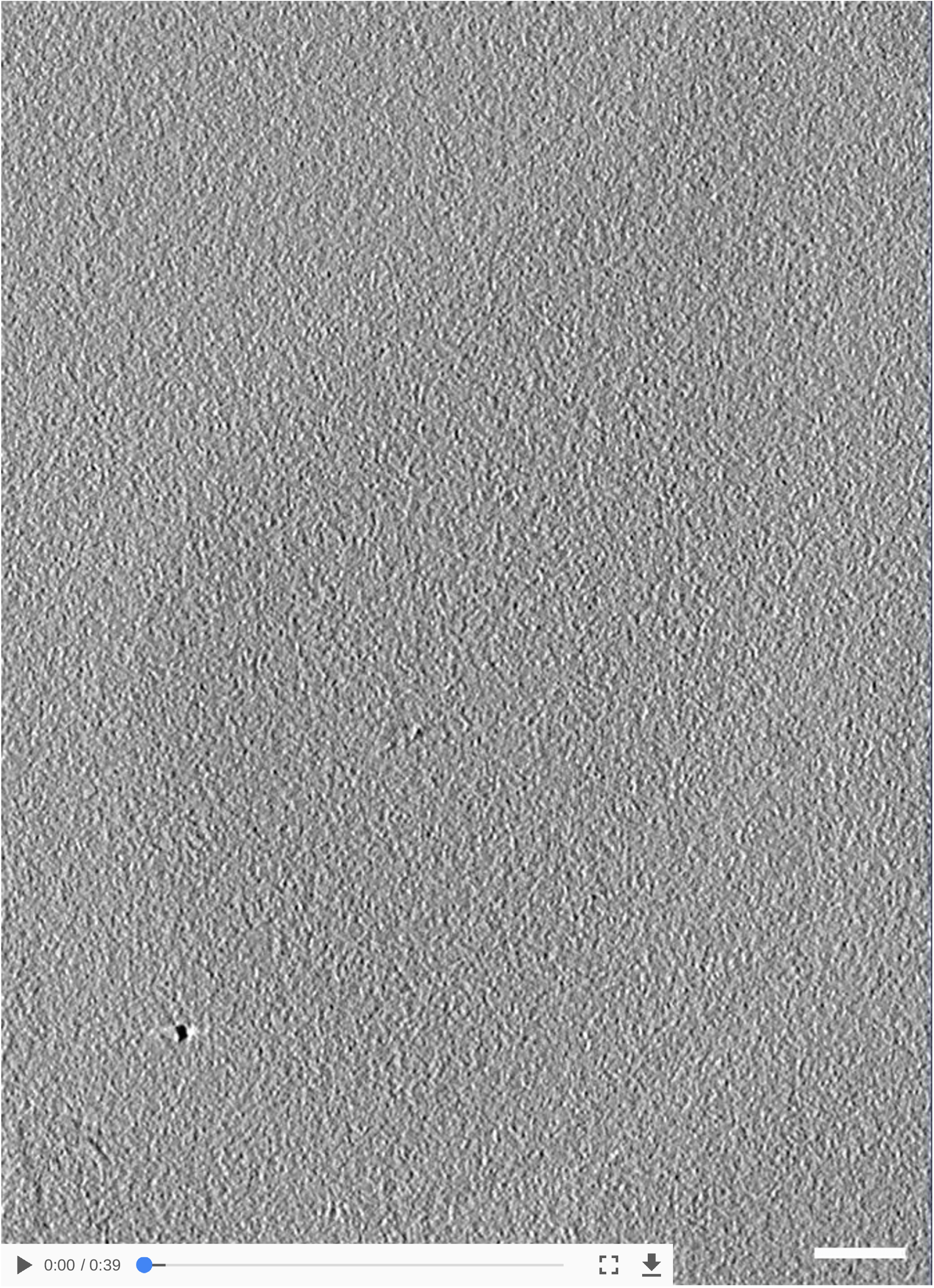
Tomogram with segmentation of WT SNAP-25 neurons. off-white= cell outline; pink = active zone; blue = synaptic vesicles; green = large vesicles; yellow = connectors, red = tethers, scale bar 100nm

### Supplementary Tables

**Table S1:** Summary of the synaptosome tomograms.

| ID | Timepoint [ms] | Vesicles per Tomogram | Tethers per AZ | AZ surface area [$\mu\text{m}^2$] | Connectors per synapse (0-250 nm) |
| --- | --- | --- | --- | --- | --- |
| Control 1 | 0 | 220 | 15 | 0.11 | 331 |
| Control 2 | 0 | 104 | 8 | 0.04 | 264 |
| Control 3 | 0 | 127 | 4 | 0.05 | 230 |
| Control 4 | 0 | 143 | 3 | 0.03 | 482 |
| Control 5 | 0 | 213 | 12 | 0.11 | 361 |
| Control 6 | 0 | 104 | 7 | 0.06 | 199 |
| Control 7 | 0 | 184 | 9 | 0.03 | 360 |
| Control 8 | 0 | 132 | 19 | 0.09 | 226 |
| Control 9 | 0 | 134 | 6 | 0.08 | 326 |
| Spray 1 | late | 697 | 5 | 0.03 | 272 |
| Spray 2 | late | 115 | 3 | 0.08 | 88 |
| Spray 3 | late | 429 | 21 | 0.19 | 882 |
| Spray 5 | early | 534 | 57 | 0.07 | 1412 |
| Spray 5_2 (second AZ, same synaptosome) | early |  | 32 | 0.15 |  |
| Spray 6 | late | 371 | 1 | 0.03 | 397 |
| Spray 7 | early | 107 | 5 | 0.02 | 156 |
| Spray 8 | late | 99 | 4 | 0.02 | 202 |
| Spray 10 | late | 76 | 4 | 0.02 | 96 |

**Table S2:** Summary of the neuron tomograms.

| ID | Mutation | Vesicles per Tomogram | Tethers per AZ | AZ surface area [$\mu\text{m}^2$] | Connectors per synapse (0-250 nm) |
| --- | --- | --- | --- | --- | --- |
| 73 | 4E | 459 | 23 | 0.23 | 269 |
| 80 | 4E | 105 | 6 | 0.21 | 84 |
| 84 | 4E | 109 | 10 | 0.12 | 107 |
| 88 | 4E | 154 | 10 | 0.13 | 159 |
| 102 | 4E | 103 | 0 | 0.29 | 94 |
| 114 | 4K | 123 | 2 | 0.09 | 68 |
| 115 | 4K | 137 | 1 | 0.26 | 70 |
| 116 | 4K | 278 | 1 | 0.13 | 154 |
| 123 | 4K | 55 | 3 | 0.09 | 52 |
| 128 | WT-KO | 243 | 12 | 0.18 | 683 |
| 132 | WT-KO | 126 | 2 | 0.24 | 110 |
| 133 | WT-KO | 505 | 27 | 0.17 | 229 |
| 134 | WT-KO | 600 | 7 | 0.34 | 144 |

## Author contributions

JR, RS, JBS, and BZ designed the study. JR, RS, and Anna K performed the experiments. KG provided access to and assistance at one of the Titan Krios microscopes. JR, RS, and BZ analyzed the data. VL, UL, and Amin K contributed to the Pyto analysis. JR, RS, JBS, and BZ wrote the manuscript with contribution from all authors. JBS and BZ supervised the project.

## Acknowledgments

We would like to thank Marek Kamínek for maintaining the electron microscope and supporting its use in Bern, and Tillmann Hanns Pape for support in specimen preparation and electron microscope operation in Copenhagen. Data was acquired on a machine supported by the [Microscopy Imaging Center (MIC) of the University of Bern](https://www.mic.unibe.ch/), a machine supported by the [Core Facility for Integrated Microscopy (CFIM) of the University of Copenhagen](https://cfim.ku.dk/publications/), and a machine supported by the Center for Cellular Imaging and NanoAnalytics, Biozentrum, University of Basel. This work was funded through the grants mentioned in the author list.

